# Airway epithelial cells and macrophages trigger IL-6-CD95/CD95L axis and mediate initial immunopathology of COVID-19

**DOI:** 10.1101/2022.08.22.504760

**Authors:** Thais F. C. Fraga-Silva, Ualter G. Cipriano, Marcilio J. Fumagalli, Giseli F. Correa, Carlos A. Fuzo, Fabiola L. A. C. Mestriner, Christiane Becari, Andrea Teixeira-Carvalho, Jordana Coelho-dos-Reis, Mayra G. Menegueti, Luiz T. M. Figueiredo, Olindo A. Martins-Filho, Marcelo Dias-Baruffi, Maria Auxiliadora-Martins, Rita Tostes, Vania L. D. Bonato

**Author notes:** Correponding author: Vania Luiza Deperon Bonato, Av Bandeirantes 3900, Ribeirao Preto, Sao Paulo 14049-900, Brazil.

## Abstract

Airway epithelial cells (AEC) are the first in contact with SARS-CoV-2 and drive the interface with macrophage to generate inflammation. To elucidate how those initial events contribute to the immunopathology or to dysregulate the immune response observed in severe and critical COVID-19, we determined the direct and indirect interactions of these cells. AEC lineage (Calu-3) infected with SARS-CoV-2 and epithelial cells (CD45^-^EpCAM^+^) from intubated COVID-19 patients showed high expression of CD95L. Infected-Calu-3 cells secreted IL-6, and expressed annexin V and caspase-3, apoptosis markers. The direct interaction of macrophages with sorted apoptotic Calu-3 cells, driven by SARS-CoV-2 infection, resulted in macrophage death and increased expression of CD95, CD95L and CD163. Macrophages exposed to tracheal aspirate supernatants from intubated COVID-19 patients or to recombinant human IL-6 exhibited decreased HLA-DR and increased CD95 and CD163 expression. IL-6 effects on macrophages were prevented by tocilizumab (anti-IL-6 receptor mAb) and Kp7-6 (CD95/CD95L antagonist). Similarly, lung inflammation and death of AEC were decreased in CD95 and IL-6 knockout mice infected with SARS-CoV-2. Our results show that the AEC-macrophage interaction via CD95/CD95L signaling is an initial key step of immunopathology of severe COVID-19 and should be considered as a therapeutic target.

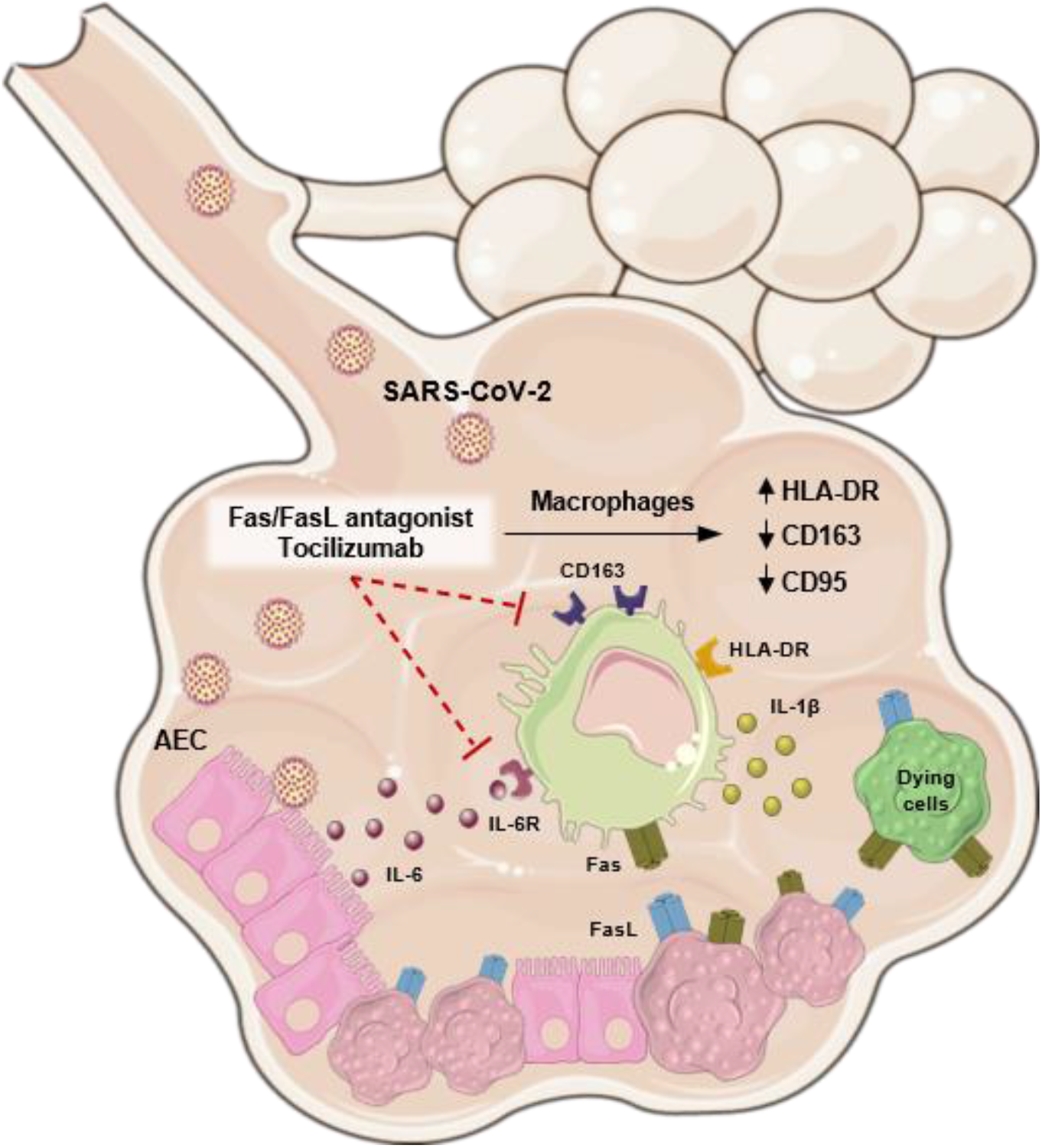

**Highlights:**

- SARS-CoV-2-infected airway epithelial cells (AEC) secrete IL-6, express Fas/FasL and undergo apoptosis;
- SARS-CoV-2-infected apoptotic AEC induces Fas/FasL expression and death in macrophages;
- IL-6 induces IL-1β secretion, reduction of HLA-DR and increase of Fas and CD163 expression in macrophages;
- Blockade of IL-6 signaling and Fas/FasL restores the expression of HLA-DR and reduces the expression of Fas and CD163, and secretion of IL-1β on isolated macrophages; in vivo, the deficiency of Fas and IL-6 decreases acute pulmonary inflammation in SARS-CoV-2-infected mice.

## Introduction

The coronavirus disease 2019 (COVID-19) pandemics started in March 2020 and it is caused by the new coronavirus SARS-CoV-2 (Severe Acute Respiratory Syndrome Coronavirus 2). SARS-Cov-2 infects primarily epithelial cells in the respiratory tract. The Spike protein on the virus surface is recognized by the angiotensin-converting enzyme 2 (ACE2), which is highly expressed in epithelial cells of the higher airways, as nasopharynx, compared to the lower airways or distal lungs (1–3). SARS-CoV-2 infection of human airway epithelial cells (AEC) *in vitro* induces a cytopathic effect (4), stimulating caspase-8-dependent IL-1β production (5), activating gene expression of inflammatory factors, and inhibiting type-I IFN production (6). Although infection of neurons, cardiovascular, renal and intestinal cells has been reported (7), SARS-CoV-2 generally exhibits a tropism for the lungs.

COVID-19 is a very complex disease, with a spectrum of clinical forms. While most of the infected subjects, approximately 80%, develop a mild disease with variable symptoms that resemble a common cold, such as cough, throat pain, fever, diarrhea, and anosmia, severe or critical patients develop acute respiratory distress syndrome (ARDS), respiratory failure and require admission to intensive care units, with O_2_ supplementation, and mechanical ventilation (8).

Even though SARS-CoV-2 particles are found in other organs, the primary cause of death is frequently respiratory failure that courses with acute bilateral pneumonia characterized by edema, capillary congestion, microthrombi, hemorrhage, vasculitis (9). Viral particles were identified within bronchial epithelial cells, alveolar type I (AT-I) and II (AT-II) cells (10), and endothelial cells (11). The infection of AEC induces disruption of intercellular junctions and loss of contact with the basement membrane while the infection of alveolar endothelial cells induces endotheliitis and endothelial injury (4,12). SARS-CoV-2 infection is also accompanied by a cytokine storm (13), which might contribute to the hemophagocytic lymphohistiocytosis that occurs in a number of viral infections (14). Patients with severe and critical forms of COVID-19 exhibit increased serum concentrations of IL-6 and IL-10, lower expression of HLA-DR on monocytes, and increased neutrophil to lymphocyte ratio in the peripheral blood (15). In addition to lymphopenia, lymphocytes obtained from blood or lungs of deceased patients express PD-1 (programmed cell death-1), an exhaustion marker (16,17). Severe COVID-19 patients are grouped in 2 immunotypes: 1-deceased patients that remain a short time at the hospital, exhibit high viral load and low diffuse alveolar damage (DAD); 2-deceased patients that remain a long time at the hospital, have lower viral load and significative DAD and immunopathology compared to the group 1 (17,18). These findings show that dysregulation of the immune response and immunopathology are hallmarks of severe COVID-19 and that the severe pulmonary disease is heterogeneous and have distinct immunotypes.

SARS-CoV-2 infects AEC, induces cell death, DAD, chemokine and cytokine secretion, acute inflammation, neutrophil influx, and neutrophil extracellular traps (NET) released during cell death (NETosis) (15,19). Postmortem examination revealed a predominance of myeloid infiltrate, mostly macrophages, along with CD4^+^ and CD8^+^ T cell migration in the pulmonary parenchyma of COVID-19 patients (7), indicating a critical role for macrophages in the immunopathology and in the dysregulation of the immune response in the lungs (20). However, little is known on the initial events that drive the interactions of lung epithelial cells and macrophages and how these early events contribute to the dysregulated immune response induced by macrophages in the lungs. Therefore, our study focused on these very early events that follows SARS-CoV-2 infection and addressed how they activate the innate response leading to hyperinflammation. We hypothesized that death of epithelial cells and the secretion of IL-6 after SARS-CoV-2 infection drive the macrophage activation, but also the macrophage dysfunction, and both contribute to the early pulmonary damage.

## Results

### SARS-CoV-2 induces CD95/CD95L expression in airway epithelial cells

SARS-CoV-2 causes a cytopathic effect in AEC, with cytokine and chemokine secretion and caspase-8-dependent IL-1β production (4,5). However, the mechanisms by which AEC trigger the innate response and inflammation are still unclear. First, we analyzed cell viability in SARS-CoV-2 (SARS2)-infected cultured Calu-3 cells, a cell line of bronchial epithelial cell, using different viral loads (MOI 0.2 and MOI 2.0). Massive cell death was detected 72 hours post-infection (**Figures 1A, 1B**) while a significant viral load (PFU/mL) was recovered from the supernatant of infected cells at 24 and 72 hours (**Figure 1C**). Since IL-6 is a critical mediator of immune dysregulation and severe respiratory failure (20), we analyzed IL-6 kinetics in SARS2-infected Calu-3 cells. SARS2 at MOI 2.0 significantly induced IL-6 secretion 6 hours post-infection compared to the MOI 0.2 and mock conditions. The peak of IL-6 secretion occurred 72 hours post-infection (**Figure 1D**). Infected Calu-3 cells secreted low levels of TNF, IL-1β and IL-10 in all conditions (data not shown).

**Figure 1.**
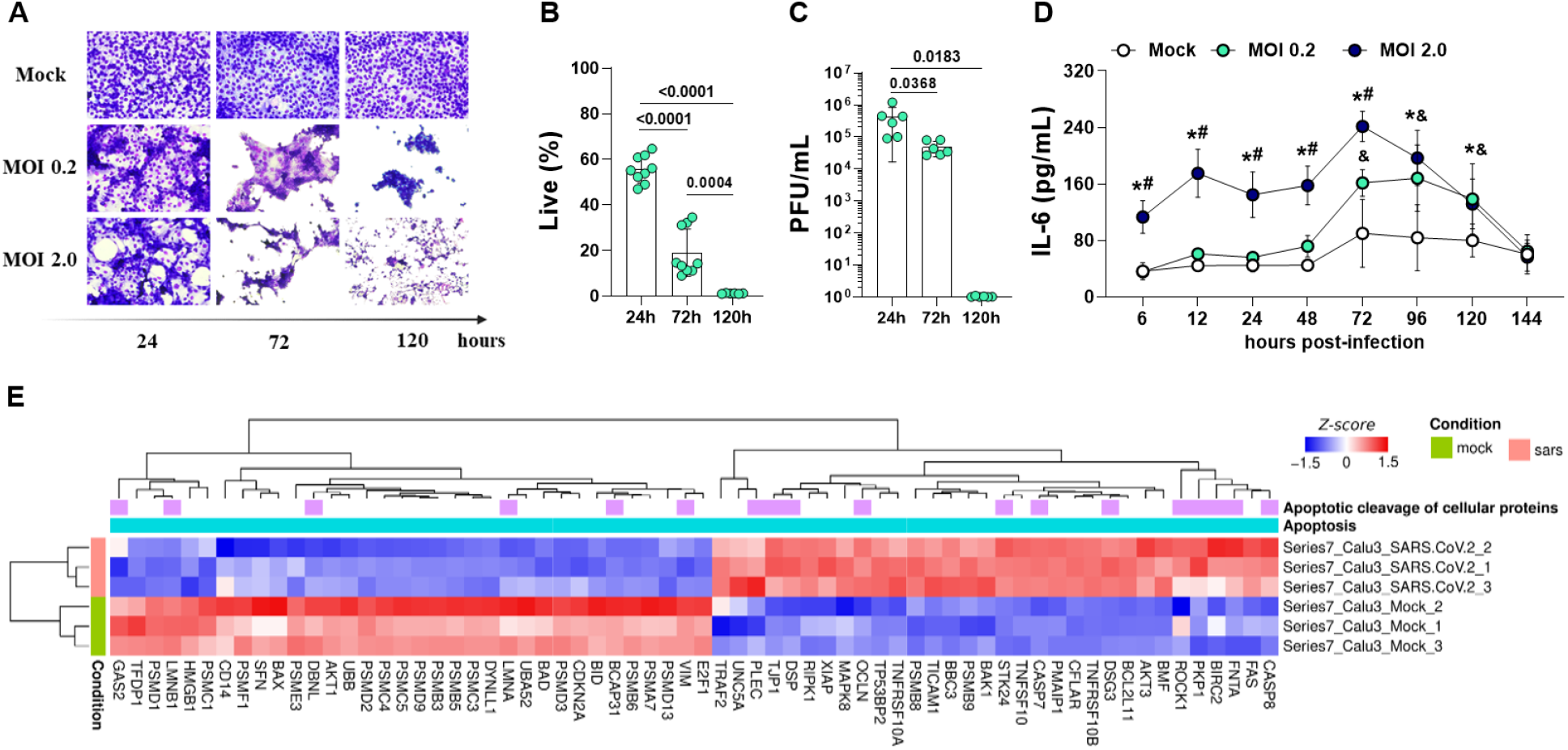
SARS-CoV-2 induces death and IL-6 secretion in airway epithelial cells. Calu-3 cells were infected with SARS-CoV-2 at multiplicity of infection (MOI) 0.2 and 2.0. Cytopathic effect was evaluated using (**A**) Panotic staining and (**B**) flow cytometric (annexin V^-^FVS^-^) analysis. (**C**) Viral load was measured by plaque-forming unit (PFU/mL) using the supernatants of Calu-3-infected cells at 24, 72 and 120 hours post-infection (p.i.) (**D**) IL-6 concentration was measured in the supernatants derived from non-infected and SARS2-infected cells at 6 to 144 hours p.i. (**E**) Normalized expression (z-score) of differentially expressed genes related to apoptosis pathways in infected and non-infected (mock) cells. Data represent the mean ± SD of at least two independent experiments performed in triplicate. (**D**) * p<0.05, MOI 2.0 *vs*. Mock; # p<0.05, MOI 2.0 *vs*. MOI 0.2, and & p<0.05, MOI 0.2 *vs*. Mock. Bars depict the p values.

Using transcriptome data obtained from Blanco-Melo et al. (6) we evaluated differentially expressed genes related to apoptotic pathways in Calu-3 cells infected with SARS2. Compared to Mock, SARS2-infected Calu-3 cells showed increased expression of pro-apoptotic genes (*CASP7, CASP8, BCL2L11*, and *CD95* – Fas receptor) and decreased expression of anti-apoptotic genes (*BIRC2*), and genes related to the proteasome complex (PSM family) or to tissue damage (*HMGB1*) (**Figure 1E**).

Most of Calu-3 cells infected with SARS2 were in late apoptotic stage (annexin V^+^FVS^+^), as confirmed by flow cytometry analysis (**Figures 2A-2D**). A low frequency of non-infected cells was also in late apoptotic stage, as those are cultured in a medium with low fetal bovine serum (**Figure 2A**). At the late infection (120h), Calu-3 cells infected with SARS2 were in necrosis compared to 24 and 72h of infection (**Figure 2D**). In addition, the expression of CD95L (Fas-ligand) and CD95, determined by CD95L or CD95 median fluorescence intensity (MFI), was significantly higher only on late apoptotic cells (**Figures 2E-2G**). Increased expression of cleaved caspase-3 in SARS2-infected Calu-3 cells (**Figure 2H**) and lower levels of HMGB1 in the supernatant of infected cells (**Figure 2I**) further support predominant cell death by apoptosis in AEC infected with SARS2.

**Figure 2.**
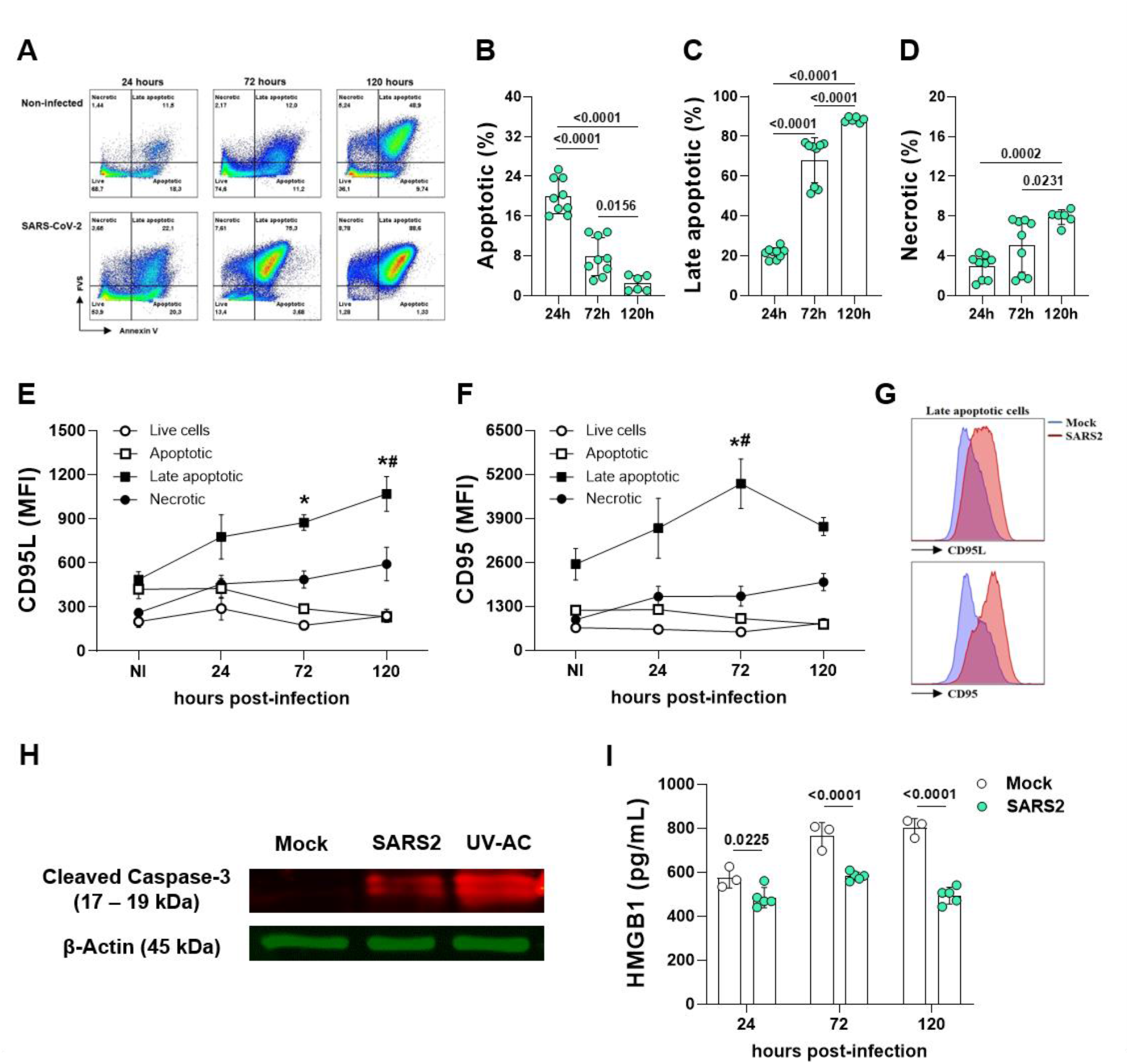
SARS-CoV-2 infection induces apoptosis and CD95/CD95L expression in airway epithelial cells. Calu-3 cells were infected with SARS-CoV-2 at MOI 0.2 and evaluated by (**A**) flow cytometry to assess the percentage of (**B**) apoptotic (annexin V^+^FVS^-^), (**C**) late apoptotic (annexin V^+^FVS^+^) and (**D**) necrotic (annexin V^-^FVS^+^) cells after 24, 72 and 120 hours p.i. The median fluorescence intensity (MFI) of (**E**) CD95L and (**F**) CD95 was determined in these cell populations and (**G**) represented in the histogram. (**H**) Representative immunoblot image of cleaved capase-3 and β-actin proteins in non-infected cells (Mock), Calu-3 cells infected with SARS-CoV-2 at MOI 0.2 for 24 hours (SARS2) and Calu-3 apoptotic cells induced by UV radiation (UV-AC, 50 mJ). (**I**) HMGB1 concentration determined in the supernatants derived from non-infected (Mock) and infected cells (SARS2) at 24, 72 and 120 hours p.i. Data represent the mean ± SD of at least two independent experiments performed in triplicate. (**E**,**F**) * p<0.05, infected *vs*. non-infected; # p<0.05, infected *vs*. 24 hours p.i. Bars depict the p values.

### Apoptotic SARS-CoV-2-infected AEC induces CD95 on macrophages

To investigate if AEC from COVID-19 patients exhibit a phenotype similar to observed in Calu-3, cell viability in 11 fresh tracheal aspirate samples collected from intubated COVID-19 patients was analyzed. Clinical and laboratorial characteristics of these patients are shown in **Table 1**. The patients, mean age was 58.9 years and 81.2% had hypertension, spent an average of 18 days in the intensive care unit (ICU), confirming severe COVID-19. Similar to the observed in Calu-3 cells, most AEC from COVID-19 patients (gated as CD45^-^EpCam^+^ cells) were dying cells (**Figure 3A**), exhibiting significant expression of CD95L (MFI), but not CD95, compared to live epithelial cells from the same sample (**Figures 3B, 3C**).

**Figure 3.**
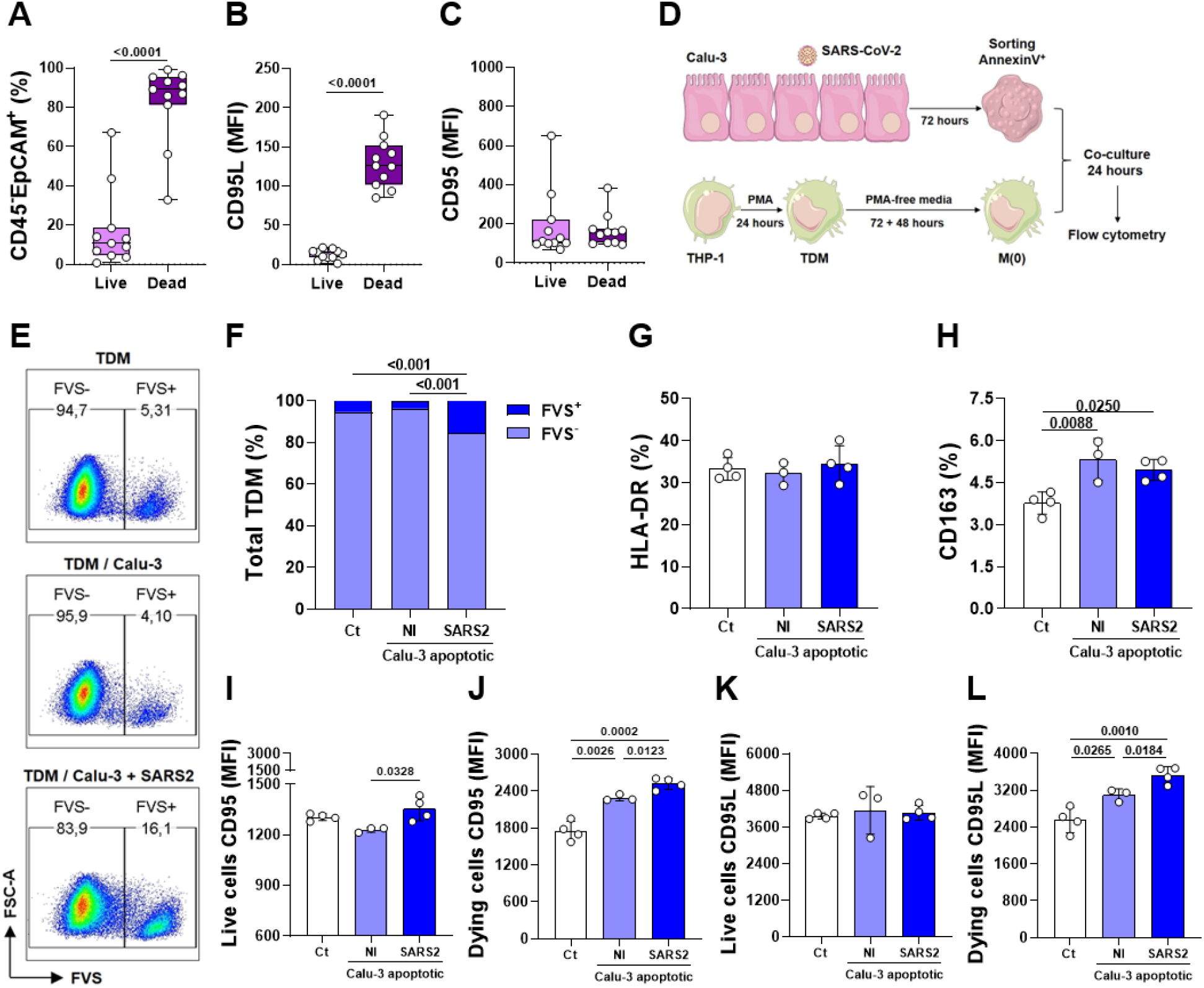
SARS-CoV-2-infected apoptotic epithelial cells induces CD95 expression and death in macrophages. Epithelial cells (CD45^-^EpCAM^+^) from tracheal aspirate of intubated COVID-19 patients were evaluated for total (**A**) live or dying cells, and the median fluorescence intensity (MFI) of (**B**) CD95L and (**C**) CD95. (**D**) THP-1-derived macrophages (TDM, 5×10^5^/mL) were co-cultured for 24 hours with sorted annexin V^+^ Calu-3 cells infected with SARS-CoV-2 (MOI 0.2) for 72 hours. (**E**,**F**) The percentage of dying cells (FVS^+^) and the percentage of (**G**) HLA-DR^+^ and (**H**) CD163^+^ in total live TDM (FVS^-^). MFI of CD95 in total (**I**) live and (**J**) dying cells and MFI of CD95L in total (**K**) live and (**L**) dying cells. Data represent the mean ± SD of one representative of two independent experiments performed at least in triplicate. Bars depict the p values.

To gain insight on how dying epithelial cells directly activate macrophages during the acute phase of COVID-19, apoptotic Calu-3 cells were sorted 72 hours post SARS2 infection and co-cultured with THP-1-derived macrophages (TDM) (**Figure 3D**). The direct contact of macrophages with SARS2-infected apoptotic Calu-3 cells, but not with non-infected apoptotic Calu-3 cells, significantly induced macrophage death (**Figures 3E, 3F**). Macrophages co-cultured with apoptotic Calu-3, SARS2-infected or non-infected, that remained alive showed no alterations in HLA-DR expression compared to control macrophage culture (**Figure 3G**). Contact with both apoptotic cells increased CD163 expression on macrophages that remained alive (**Figure 3H**). Increased expression of CD95 was observed in live and dying macrophages (**Figure 3I, 3J**), and increased expression of CD95L was observed only in dying cells (**Figures 3K, 3L**). Therefore, the direct interaction of SARS2-infected apoptotic AEC with macrophages resulted in death of these innate leukocytes and positive regulation of CD95, mainly in dying macrophages.

### IL-6 induces macrophage activation and dysfunction

Considering that SARS2-infected Calu-3 cells secrete IL-6 (**Figure 1D**), and IL-6 is associated with progression, severity and mortality in COVID-19 (21–23), the indirect (IL-6-mediated) effect of epithelial cells on macrophages was also investigated. The secretion of chemokines and cytokines was first determined in 28 samples of frozen tracheal aspirates collected from intubated COVID-19 patients. Clinical and laboratorial characteristics of patients are shown in **Table 2**. The 28 severe patients, mean age was 65.29 years and 81.8% had hypertension, spent an average of 21.6 days in the ICU. IL-1β, IL-6 and TNF were the cytokines with the highest concentrations in the supernatants of tracheal aspirate from these COVID-19 patients. Among those, IL-6 was detected in higher concentrations. Samples from the 28 intubated patients were divided in low (17 patients) and high (11 patients) IL-6-producers (**Figure 4A**). High producers showed increased levels of HMGB1 and ATP in tracheal aspirate samples compared with the low IL-6-producers (**Figures 4B, 4C**). Despite the increased levels of some mediators, no differences were observed in CD95L expression on AEC cells eluted from frozen tracheal aspirate samples (data not shown) or in the outcome of the disease (**Table 2**) between low and high IL-6-producers. However, CD95L was associated with a greater incidence of death, and high IL-6-producers exhibited a significant correlation with TNF and HMGB1 (**Figure 4D**).

**Figure 4.**
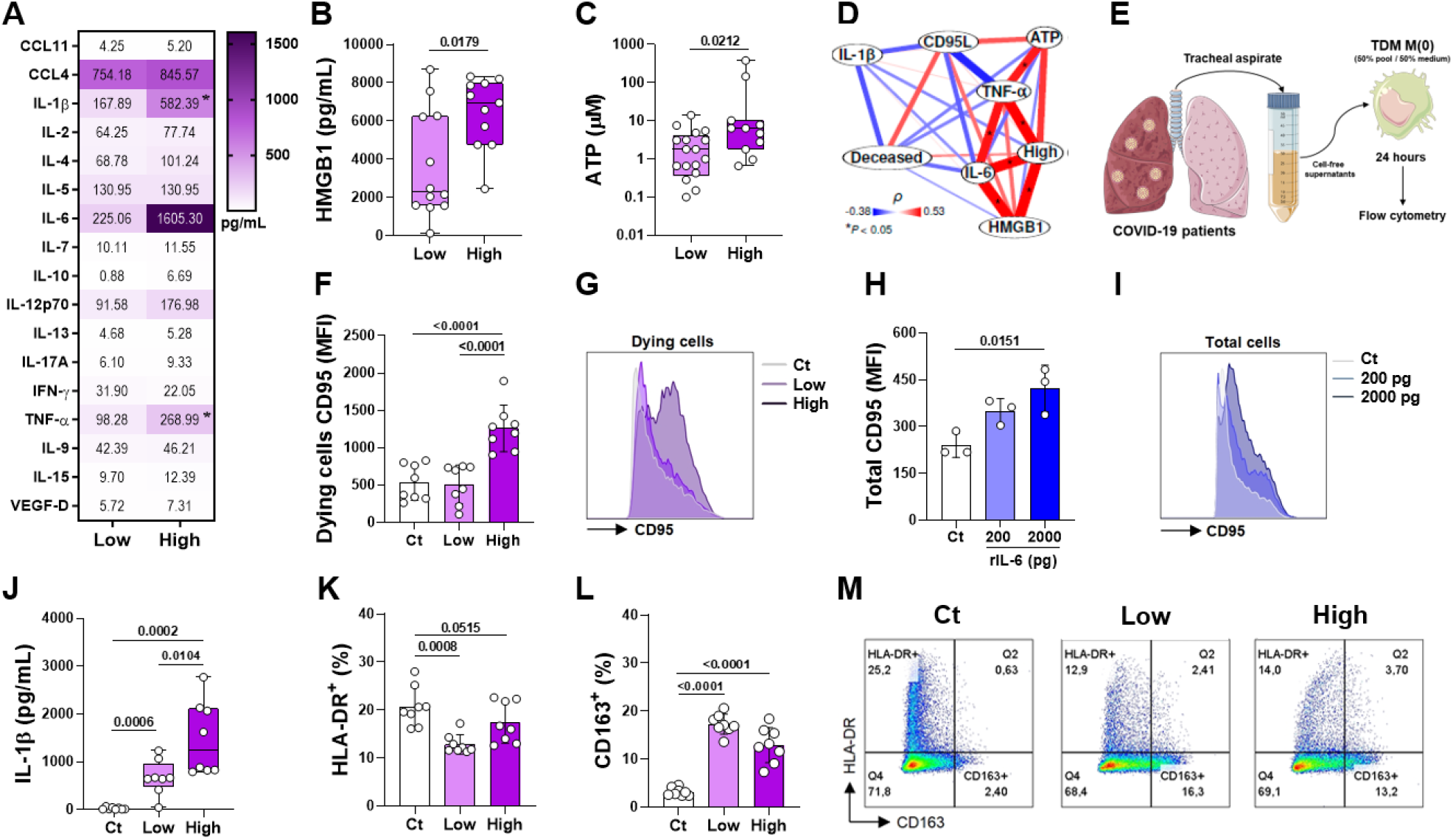
High IL-6 levels induce CD95 expression and IL-1β production in macrophages. (**A**) Inflammatory mediators were determined in tracheal aspirate samples obtained from intubated COVID-19 patients, who were sorted in pools of low and high IL-6-producers. Concentration of (**B**) HMGB1 and (**C**) ATP were measured in the cell-free supernatants derived from tracheal aspirate samples. (**D**) Correlogram of inflammatory mediators and clinical outcomes in high IL-6-producers. (**E**) THP-1-derived macrophages (TDM, 5×10^5^/mL) were cultured for 24 hours with low and high IL-6 cell-free supernatants. The median fluorescence intensity (MFI) of CD95 was assessed in (**F**,**G**) dying TDM (FVS^+^) cultured with tracheal aspirate supernatants and in (**H**,**I**) total TDM cultured with low (200 pg) and high (2000 pg) concentrations of human recombinant IL-6. IL-1β detection in the supernatants (**J**) and percentage of HLA-DR (**K**,**M**) and CD163 (**L**,**M**) evaluated in total live TDM (FVS^-^) cultured with tracheal aspirate supernatants. Data represent the mean ± SD of two independent experiments performed in quadruplicate for tracheal aspirate cultures and one experiment for recombinant human IL-6 cultures. (**A**) * p<0.05, low *vs*. high; (**D**) * p<0.05, Spearman’s correlation. Bars depict the p values.

Pools of cell-free supernatants of frozen tracheal aspirate derived from low and high IL-6-producers were sorted to evaluate the effect of the inflammatory milieu on steady-state macrophages. TDM were cultured with supernatants of low or high producers of IL-6 for 24 hours (**Figure 4E**). TDM cultured with supernatants of high IL-6-producers showed higher expression of CD95 (**Figures 4F, 4G**) with no differences in the expression of CD95L (data not shown). IL-6 concentrations detected in the supernatants of tracheal aspirates were very variable. The range of IL-6 in the supernatants of tracheal aspirate varied from 6.43 – 529.25 pg/mL for low producers and from 767.31 – 1,792.87 pg/mL for high producers. As the mean values from lower and higher samples evaluated were 225.06 and 1,605.30 pg/mL, cultures of TDM were stimulated with human recombinant IL-6 (rIL-6) at 200 pg and 2.000 pg to mimic low and high concentrations of IL-6. CD95 expression was greater in macrophages exposed to the higher rIL-6 concentrations (**Figures 4H, 4I**), similarly to the observed with supernatants derived from tracheal aspirate samples.

Both supernatants, high and low IL-6-producers induced IL-1β by macrophages. High IL-6-producers induced a significant augment in the concentrations of IL-1β compared to stimulation in the presence of low IL-6-producers (**Figure 4J**). Of note, the concentrations of IL-1β in the supernatants of TDM cultures were determined subtracting the concentrations previously detected in supernatants of the tracheal aspirate. Both supernatants (high and low producers) induced decreased percentage of HLA-DR^+^ (**Figures 4K, 4M**) and increased percentage of CD163^+^ TDM (**Figures 4L, 4M**).

### Blockade of the IL-6-CD95/CD95L axis augments HLA-DR and reduces CD163 on macrophages

To confirm that the IL-6-CD95/CD95L axis leads to the death of AEC and to both macrophage activation and dysfunction, the effects of Kp7-6 (CD95/CD95L antagonist) and tocilizumab (IL-6 receptor antagonist) were determined. Calu-3 cells treated with Kp7-6 for 2 hours and then infected with SARS2 (MOI 0.2) for 24 hours showed significant death reduction compared with vehicle-treated infected cells (**Figure 5A**).

**Figure 5.**
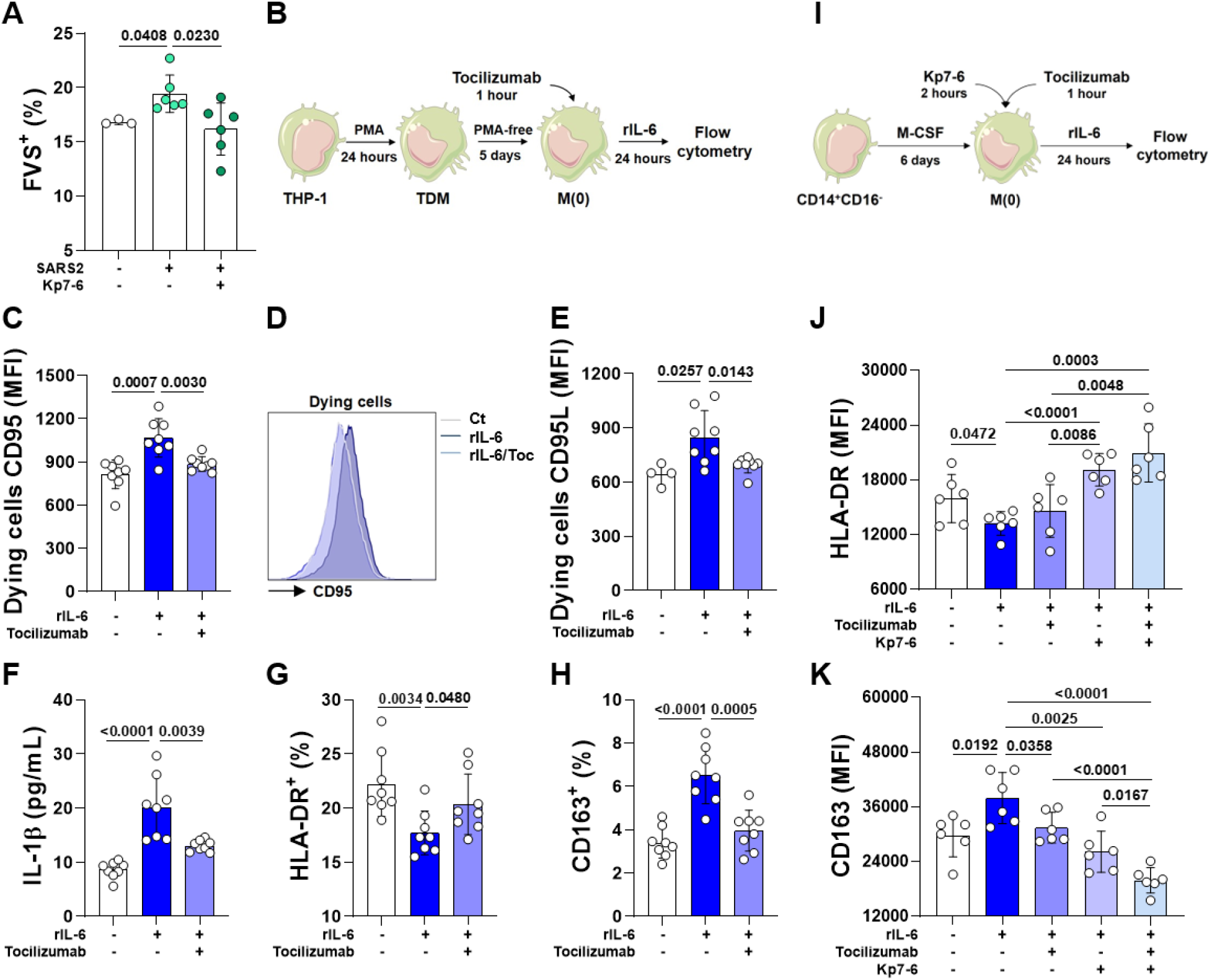
Blockade of IL-6 and CD95 prevents macrophage dysfunction induced by IL-6. Calu-3 cells were infected with SARS-CoV-2 at MOI 0.2 and evaluated by flow cytometry. (**A**) Percentage of total dying cells (FVS^+^) 24 hours p.i. (**B**) THP-1-derived macrophages (TDM, 5×10^5^/mL) were cultured for 24 hours with recombinant human IL-6 (4000 pg/mL) after pre-treatment with tocilizumab (100 µg/mL) for 1 hour and evaluated by flow cytometry. The median fluorescence intensity (MFI) of (**C**,**D**) CD95 and (**E**) CD95L in total dying TDM (FVS^+^). (**F**) IL-1β detection in the supernatants and percentage of (**G**) HLA-DR^+^ and (**H**) CD163^+^ cells evaluated in total live TDM (FVS^-^). (**I**) Monocyte derived macrophages (MDM, CD14^+^CD16^-^, 5×10^5^/mL) were cultured for 24 hours with recombinant human IL-6 (4000 pg/mL) after pre-treatment with Kp7-6 (100 µg/mL) for 2 hours and/or with tocilizumab (100 µg/mL) for 1 hour. Expression of HLA-DR (**J**) and CD163 (**K**) were evaluated by flow cytometry in total live human macrophages (FVS^-^). Data represent the mean ± SD of two independent experiments performed in quadruplicate (TDM) or triplicate (MDM). Bars depict the p values.

TDM were also treated with tocilizumab 1 hour before rIL-6 stimulation (**Figure 5B**). rIL-6 increased CD95 and CD95L expression (**Figures 5C-5E**), induced the production of IL-1β (**Figure 5F**), decreased the expression of HLA-DR (**Figure 5G**) and increased the expression of CD163 on macrophages (**Figure 5H**). The pre-treatment with tocilizumab inhibited rIL-6 effects on TDM (**Figures 5C-5H**).

Next, monocyte derived macrophages (MDM) were pre-treated with tocilizumab or Kp7-6 and stimulated with rIL-6 (**Figure 5I**). No changes were found in CD95 expression and IL-1β was not detected in the supernatants of MDM cultures (data not shown). However, MDM stimulated with rIL-6 exhibited results similar to those described for TDM: low HLA-DR expression (**Figure 5J**) and high CD163 expression (**Figure 5K**). Tocilizumab alone partially reverted the effect of rIL-6 and reduced CD163 expression, while Kp7-6 showed a more pronounced effect, with increased HLA-DR expression and decreased CD163 expression. The combined treatment, tocilizumab plus Kp6-7, produced an additional effect on the increased expression of HLA-DR and on the decreased expression of CD163 (**Figures 5J, 5K**).

### Up regulation of MHCII on alveolar macrophages in infected CD95^-/-^ but not in WT mice

To investigate the role of IL-6-CD95/CD95L axis *in vivo*, we used a mouse model to evaluate acute pulmonary inflammation. Although SARS-CoV-2 infection of C57BL/6 Wild Type (WT) mice is not progressive compared to humanized ACE2 mouse, WT mouse infection is used to evaluate the immunopathology (24). An intense inflammatory infiltration was observed after 3 days of infection (**Figures 6A-6C**) in the lungs of WT mice. CD95 (lpr^-/-^) and il-6 (il-6^-/-^) knockout mice showed moderate pulmonary inflammation in the lungs compared to infected WT mice (**Figure 6A-6C**).

**Figure 6.**
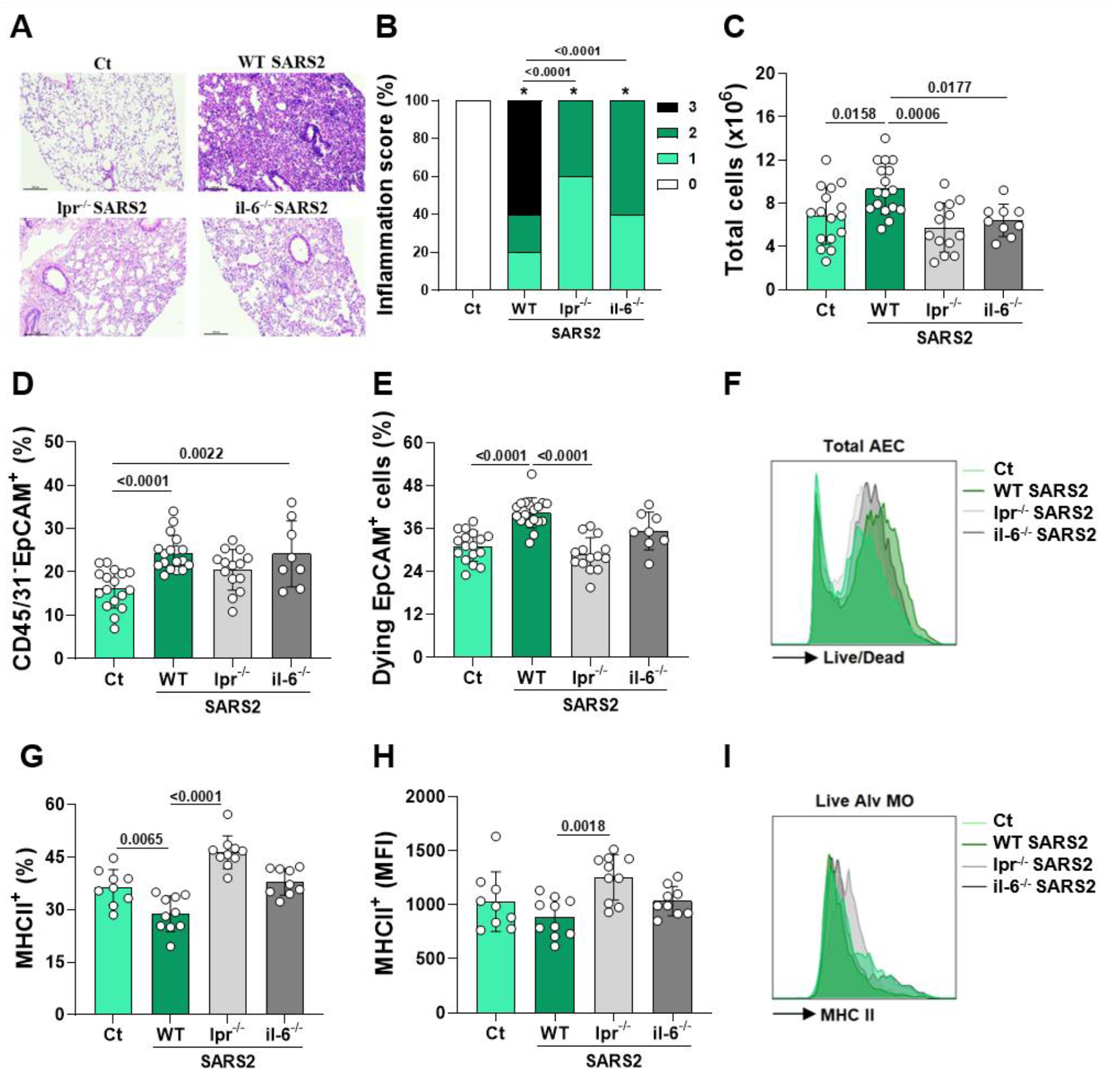
Lack of CD95 and IL-6 signaling reduces SARS-CoV-2-induced lung inflammation in mice. C57BL/6 wild type (WT) and CD95 (lpr^-/-^) and IL-6 (il-6^-/-^) knockout mice were intranasally infected with SARS-CoV-2 (SARS2, 5×10^5^ PFU) and evaluated 3 days post-infection. (**A**) Representative histopathological analysis, (**B**) score of inflammatory infiltration and (**C**) total cells count in the lungs. The percentage of (**D**) total AEC cells (CD45/31^-^EpCAM^+^) and (**E**,**F**) dying AEC cells (CD45/31^-^EpCAM^+^Live/Dead^+^) were determined by flow cytometry. Live alveolar macrophages (Alv MO - L/D^-^CD11b^-^CD11c^+^ SiglecF^+^) were evaluated to assess (**G**) the percentage of MHCII^+^ and (**H**,**I**) the median fluorescence intensity (MFI) of MHCII by flow cytometry. Data represent the mean ± SD of two - four independent experiments (n = 8-16/group). Bars depict the p values.

SARS2 infection increased the percentage of epithelial cells (AEC - CD45^-^CD31^-^EpCAM^+^) in the lungs of WT and il-6^-/-^ mice compared to non-infected mice (Ct group), while infected lpr^-/-^ group showed no difference compared to Ct group (**Figure 6D**). However, lpr^-/-^ infected mice showed a significant lower percentage of lung dying AEC, which was significantly higher in the infected WT group (**Figures 6E, 6F**). SARS2 infection decreased the percentage of alveolar macrophages (CD11b^-^CD11c^+^SiglecF^+^ cells) that expressed MHCII in the lungs of WT mice, but an increase of both percentage and expression of MHCII (MFI) was observed in the lungs of lpr^-/-^ mice (**Figures 6G-6I**). These findings show that CD95/CD95L signaling induces AEC death and decreases MHCII expression in alveolar macrophages.

## Discussion

The initial steps that orchestrate the activation of macrophages, the most abundant leukocyte in the lungs of deceased COVID-19 patients (7), are still unknown. Using various experimental approaches, we tested the hypothesis that the death of epithelial cells triggers the inflammatory response of macrophages and generates immunopathology. In the present study we showed that apoptosis of AEC mediated by activation of the CD95/CD95L axis after SARS-CoV-2 infection induces both macrophage activation (production of IL-1β and expression of CD95) and macrophage dysfunction (reduction of HLA-DR and increase of CD163). However, the macrophage dysfunction requires also the participation of CD95, which is up regulated, directly and indirectly, via contact of macrophages with epithelial cells and stimulation of macrophages by IL-6, respectively. CD95 signaling increases the production of IL-1β, up regulates the expression of CD163 and down regulates HLA-DR (MHCII) on macrophages, setting the initial steps for the interactions between epithelial cells and macrophages. Our study shows a CD95 non-canonical pathway that induces inflammatory response and macrophage dysfunction, and triggers acute immunopathology in COVID-19.

The cytopathic effect of SARS-CoV-2 is mainly associated with apoptosis of human airway epithelial cells, cell fusion, destruction of epithelium integrity and cilium shrinking (25,26). Furthermore, apoptosis was described in AT-I and AT-II cells in lung sections from postmortem COVID-19 patients and also in lung tissue from non-human primates infected with SARS-CoV-2 (27). SARS-CoV-2-induced apoptosis is mediated by caspase-8 activation (5) and associated with viroporin ORF3a, a conserved coronavirus accessory protein (28). Here we show that Calu-3 cells infected with SARS-CoV-2 exhibit increased expression of annexin V, CD95 and CD95L on the cell surface and increased cleaved/active caspase-3. Furthermore, we show that most epithelial cells, obtained from severe COVID-19 patients, i.e. airway epithelial cells eluted from fresh tracheal aspirate, are dying and present a significant high expression of CD95L, confirming the *in vitro* findings.

RNA viruses, such as influenza A and respiratory syncytial virus (RSV), activate CD95 (Fas) gene expression and induce apoptosis of infected cells, which favors/guarantee efficient viral egress (29,30). Influenza virus also induces co-expression of Fas and FasL on infected cells, triggering apoptosis when the infected cells come into contact with each other (31). These studies show that CD95/CD95L pathway might be a viral egress pathway. Our results show a similar effect of SARS-CoV-2 on Calu-3 cells and in AEC obtained from tracheal aspirates of severe COVID-19 patients. Treatment of Calu-3 cells with CD95/CD95L antagonist decreased the number of dead cells to levels seen in untreated, non-infected cells, reinforcing that CD95/CD95L on epithelial cells is key for the process. This is an important finding considering that critical COVID-19 patients with higher viral load remained in the ICU for shorter periods and died before critical patients in the ICU with lower viral load (18,32). However, the pro-inflammatory role of CD95/CD95L (Fas/FasL) pathway was also described in bronchiolar epithelial cells and resulted in CXCL8 release, lung injury and pulmonary fibrosis (33). The cytokines and chemokines released by apoptotic cells, including IL-6, CXCL8, CXCL1, CCL2, and GM-CSF, act as a “find me” signal for apoptotic cells (34) and IL-6 induction upon CD95 (Fas) ligation has been described in a dose- and time-dependent manner (35). Here we show that CD95/CD95L (Fas/FasL) pathway contributes for IL-1β released by apoptotic cells, augment of CD163 and decrease of HLA-DR on macrophages. Indeed, when we infected lpr (CD95) knockout mice, we observed less death of AEC and increase of MHCII on alveolar macrophages, confirming our hypothesis and reinforcing a role for the CD95/CD95L pathway on immunopathology.

Similarly, our results show that the peak production of IL-6 coincided with the peak of CD95 expression, while the expression of CD95L remained high in infected Calu-3 cells. In addition to the frequency of dying AEC, we also found increased levels of ATP, HMGB-1, IL-6 and IL-1β in tracheal aspirates of critical COVID-19 patients. Although a small proportion of AEC were in necrosis (annexin V^+^FVS^+^), it is unlikely that they are the source of ATP, HMGB-1 and IL-1β. These critical patients were divided in high and low responders based on IL-6 production, considering that IL-6 is associated with progression, severe forms, and high mortality of disease (36). Greater IL-6 levels (obtained from supernatants from higher responders’ patients) are linked to increased IL-1β release by macrophages. IL-1β secretion was dependent on the concentration of IL-6 in the milieu, and induced macrophage dysfunction, characterized by increased CD163 and decreased HLA-DR expression. Assuming that AEC is the primary source of IL-6 in the lungs, we used recombinant IL-6 and showed that, similar to the observed in tracheal aspirate supernatants, IL-6 stimulated the secretion of IL-1β and the expression of CD163, and decreased the expression of HLA-DR, effects that were prevented by the treatment with tocilizumab and/or with Kp7-6 (CD95/CD95L antagonist). In a similar way, IL-6 concentrations in the blood are inversely associated with HLA-DR expression on CD14^+^ monocytes of COVID-19 patients and the addition of tocilizumab in the plasma-enriched medium of cells partially restores the expression of HLA-DR (20).

IL-6 also positively regulates the expression of CD95 on macrophages, being the expression of CD95 higher on macrophages stimulated with supernatants from high IL-6 responders. In addition, the apoptosis of SARS-CoV-2-infected AEC, but not apoptosis of non-infected AEC, induced CD95 expression on live macrophages. These results indicate that the activation of macrophages by AEC depends on CD95 and occurs directly and indirectly: through the contact between AEC and macrophages via CD95L and CD95, respectively; and by IL-6, secreted by epithelial cells that up regulates CD95 on macrophages. Although described as a death receptor, CD95 may mediate non-apoptotic activities that involve cell proliferation, activation and cytokine secretion (37,38). Cytokine secretion in MDM and in a macrophage cell lineage (RAW264.7) is independent of caspase, but relies on MyD88 and NF-κB signaling, suggesting that CD95-activated macrophages initiate and perpetuate acute inflammation and tissue injury (39,40).

Our findings show that recombinant IL-6 (or high levels of IL-6 from tracheal aspirates) positively regulates CD95 and CD163 expression, and reduces the expression of HLA-DR (MHCII) on macrophages. Aberrant macrophage activation described in the lungs of deceased COVID-19 patients suggests that myeloid cells are a major source of dysregulated inflammation in COVID-19 (20,41). IL-6 is an important cytokine involved in the immunopathogenesis of COVID-19 and blockade of IL-6 signaling is considered an emerging approach (42,43). Tocilizumab has been evaluated in at least 27 current clinical trials in COVID-19 (43), and is linked to improved clinical signs, reduced time and need for ventilation and, decreased mortality (44-46). Here we show that CD95/CD95L antagonist played an additional effect on tocilizumab action preventing IL-6/IL6R signaling. Furthermore, using a mouse model of immunopathology, we show that CD95 exhibited a more evident role in SARS-CoV-2-induced acute lung inflammation compared to IL-6.

Our results add new pieces to the complex scene of COVID-19 disease, especially in the initial steps that trigger the inflammatory response with focus on the role of epithelial cell and its interface with macrophages in the generation of the immunopathology. Finally, our results elucidate the function of the IL-6-CD95/CD95L axis in the interaction of SARS-CoV-2-infected AEC with macrophages and their roles in generating immunopathology, revealing CD95/CD95L as new target to be discussed as immunotherapy for the early time of severe/critical COVID-19.

## Materials and Methods

### Ethics statement

This study was approved by the local Ethic Committee (CAAE: 30816620.0.0000.5440). All participants or family members have provided informed consent. Six weeks old female C57BL/6 mice were used for SARS-CoV-2 infection of the Clinics Hospital of Ribeirao Preto of the University of Sao Paulo (SP, Brazil). Animals were obtained from the breeding facility of Ribeirao Preto Medical School, University of Sao Paulo (Ribeirao Preto, SP, Brazil). All animals were maintained in sterile environmental conditions in a ventilated rack (Alesco, Monte Mor, SP, Brazil) and received sterile food and water. All experiments were performed according to the local Ethics Committee on Animal Experimentation (Protocol Number 114/2021).

### Study participants and sample collection

The COVID-19 group included 39 patients who were admitted to the Intensive Care Unit (ICU) of the Clinics Hospital of Ribeirao Preto of the University of Sao Paulo and need mechanical ventilation. COVID-19 infection was confirmed by a positive RT-PCR test. Tracheal aspirate samples (2-5 mL) were collected by aspiration into sterile tracheal secretion collectors in the early morning routine visit of patients for airway cleansing and only productive secretion was used in this study. Tracheal aspirate samples were manipulated in a biosafety level 3 laboratory (Department of Biochemistry and Immunology, Ribeirao Preto Medical School, University of Sao Paulo, Ribeirao Preto, Sao Paulo, Brazil) in at least 2 hours post collection. All sample were diluted in 1:2 in phosphate buffered saline (PBS) and filtered through cell strainers (70 µm) into polypropylene tubes to remove mucous clumps and cell aggregates. Samples were further centrifuged at 24,000 x g and filtered through 0.45 µm syringe filter to assess cell-free supernatants.

### SARS-CoV-2

The WT SARS-CoV-2 lineage (Genbank access MT126808.1) which are isolated from clinical samples in Brazil (51). Viral stocks were propagated in African green monkey kidney epithelial (Vero E6) cells (ATCC CRL-1586) cultured with Dulbecco’s Modified Eagle’s Medium (DMEM, Gibco) supplemented with 1% penicillin/streptomycin and 2% Fetal Bovine Serum (FBS, Gibco) and maintained at 37°C and 5% CO2. Viral titration was performed by plaque assay as previous described (47). For viral RNA quantification, RNA was extracted with TRIzol (Thermo, USA) reagent with 20 mg of lung homogenate, according to manufacturer’s recommendation and qRT-PCR was performed using TaqMan Fast Virus 1-Step Mix (Applied Biosystems, USA) according to the manufacturer’s recommendations. SARS-CoV-2 primers and probe were designed to target a 100bp region from RNA-dependent-RNA polymerase gene (RdRp) of all three SARS-CoV-2 variants (Forward: 5’ – GTGAAATGGTCATGTGTGGCGG – 3’; Reverse: 5’ – CAAATGTTAAAAACACTATTAGCATA – 3’ and Probe: ‘5-FAM – CAGGTGGAACCTCATCAGGAGATGC – BHQ1-3’) and the reactions were performed using a StepOnePlus Real-Time PCR system (Applied Biossystems), according (24).

### Cell culture

Human adenocarcinomic lung epithelial (Calu-3, BCRJ 0264) and human peripheral blood monocyte (THP-1, BCRJ 0234) cell lines were maintained at 37ºC and 5% CO2 in DMEM (1 g/L D-glucose, GIBCO) supplemented with 20% fetal bovine serum (FBS, Gibco) and 1% penicillin/streptomycin (Gibco). Calu-3 infection with SARS-CoV-2 was performed at an MOI of 0.2 or 2.0 in DMEM (4.5 g/L D-glucose, GIBCO) supplemented with 2% FBS, 1% penicillin/streptomycin, 4 mM L-glutamine, 10 mM non-essential amino acids and 1 mM sodium pyruvate, as previously described (6). THP-1 cells were differentiated into a macrophage-like phenotype (TDM, THP-1-derived macrophages) using phorbol-12-myristate-13-acetate (PMA, 5 ng/mL) for 24 hours and 5 days for rest period in PMA-free medium, as described by Baxter et al. (48). Then, TDM was co-cultured with apoptotic Calu-3, stimulated with 50% cell-free supernatants derived from tracheal aspirate of patients or stimulated with human recombinant IL-6. Human monocytes were differentiated into a macrophage-like phenotype (MDM, monocyte derived macrophages) using M-CSF (50 ng/mL, Miltenyi Biotec, Bergisch Gladbach, Germany) for 6 days at 37ºC and 5% CO_2_ in Iscove’s Modified Dulbecco’s Medium (IMDM, Lonza) supplemented with 5% FBS and 1% penicillin/streptomycin, adapted from Hoepel et al. (49). Monocytes (CD14^+^CD16^-^) were sorted by columns with magnetic beads (Miltenyi Biotec) from total mononuclear cells isolated by Ficoll gradient from the peripheral blood of healthy volunteers. MDM were then stimulated with recombinant human IL-6.

### AEC kinetic and co-culture

For kinetic experiments, Calu-3 were infected with SARS-CoV-2 at an MOI of 0.2 or 2.0 for 6, 12, 24, 48, 72, 96, 120 and 144 hours at 37ºC and 5% CO_2_. Viral load and inflammatory mediators were assessed in supernatants, protein was evaluated in lysate cells and the type of cell death was evaluated in detached cells after treatment with trypsin (Gibco). For treatment experiment, approximately 5×10^5^ Calu-3 cells were pre-treated with kp7-6 (100 ug/mL, CD95/CD95L antagonist, Merck) for 2 hour and then infected with SARS-CoV-2 at an MOI of 0.2. For co-culture experiments, Calu-3 were infected with SARS-CoV-2 at an MOI of 0.2 for 72 hours and detached cells were stained with FVS and annexin V. Apoptotic cells (annexin V^+^) were freshly sorted in FACS Melody (BD Biosciences, San Jose, CA) allocated in a biosecurity level 3 laboratory (Department of Biochemistry and Immunology, Ribeirao Preto Medical School, University of Sao Paulo, Ribeirao Preto, Sao Paulo, Brazil). Sorted AEC were then co-cultured with TDM for 24 hours in an equal proportion (1 apoptotic cell : 1 macrophage).

### Transcriptome analysis

The transcript expression data was obtained from public data deposited at *Gene Expression Omnibus* database (50). We selected count raw data for the transcript reads for independent biological triplicates for SARS-CoV-2 (USA-WA1/2020 strain) infected and mock Calu-3 deposited by Blanco-Melo (6) with id GSE147507. Differential gene expression analysis of SARS-CoV-2 infected versus mock control was carried out using with *DESeq2*. The *p*-values derived from differential expression analysis were adjusted (*p*_adj_) for multiple testing using the Benjamini and Hochberg method (51). Differentially expressed gene (DEG) signature was defined using a threshold for absolute value of fold-change at log_2_ basis > 0.5 and *p*_adj_ < 0.05. The DEG list was submitted to over-representation analysis using *cluster Profiler* over *Reactome pathways* annotations within BH adjusted *p*-value < 0.05. The heatmap for normalized gene expression was constructed using *pheatmap*. The data was analyzed in *R* environment (https://www.R-project.org/).

### Macrophage stimulation and treatments

Approximately 2.5×10^5^ TDM were stimulated with a pool of cell-free supernatants derived from TA samples of COVID-19 patients in an equal proportion with culture medium (1:1 v/v) for 24 hours at 37ºC and 5% CO_2_. Approximately 2.5×10^5^ TDM and MDM were stimulated with recombinant human IL-6 (200 or 2000 pg, Peprotech) for 24 hours at 37ºC and 5% CO_2_. Approximately 2.5×10^5^ TDM cells were pre-treated with tocilizumab (100 µg/mL, Actemra) for 1 hour and then stimulated with recombinant IL-6. Approximately 5×10^5^ MDM were pre-treated with kp7-6 (100 µg/mL) for 2 hours and pre-treated with tocilizumab (100 µg/mL) for 1 hour and then stimulated with recombinant IL-6. Inflammatory mediators were assessed in supernatants and protein expression was analyzed using flow cytometry in detached cells after treatment with EDTA (5 mM).

### Mice infection and sample collection

Mice were infected with 5×10^5^ PFU of WT SARS-CoV-2 variants by intranasal route at previously described (24). Mice were euthanized 3 days post infection and the lungs were collected for analysis. The superior right lobe was gently perfused with 10% formaldehyde solution, embedded in paraffin, sectioned at 4 µm thickness and stained with Harris’s hematoxylin and eosin (H&E) for histopathological analysis. The left lung was collected in complete RPMI media and processed for flow cytometry analysis. The middle and inferior right lobes were weighted and homogenized with PBS (1:5 w/v) using a 5 mm stainless steel bead (Qiagen, USA) and a TissueLyser LT (Qiagen, USA), (50 Hz for 5 min), centrifuged at 10,000 x g for 10 min, and the supernatant was collected and used for viral load measurement by plaque assay and qRT-PCR and cytokines quantification.

### Soluble immune mediators

IL-1β, IL-6, IL-10, and TNF levels were determined in the supernatants of Calu-3 and macrophage cultures using ELISA kits following the manufacturer’s instructions (R&D Systems). HMGB-1 was determined in the cell-free TA supernatants and in the supernatants of Calu-3 cultures using immunoassay following the manufacturer’s instructions (Novus Biologicals). Adenosine 5′-triphosphate (ATP) was assessed in the cell-free TA supernatants using bioluminescent assay kit following the manufacturer’s instructions (Sigma-Aldrich). The levels of soluble immune mediators were measured in TA samples using a high-throughput microbeads array (Bio-Plex Pro™ Human Cytokine 27-plex Assay, Bio-Rad Laboratories, Hercules, CA, USA), following the manufacturer’s instructions. The results were expressed in pg/mL according to standard curves for each immune mediator using a fifth parameter logistic fit analysis. Cytokine profile in the lung of infected mice was determined using the Cytometric Bead Array (CBA) Mouse Th1/Th2/Th17 Cytokine kit (BD Biosciences) according to manufacturer instructions and samples were acquired on BD FACSCanto (BD Biosciences, USA) and analyzed using FCAP Array v3.0 (BD Biosciences, USA).

### Flow Cytometry

Calu-3 were detached with trypsin (Gibco) and TDM or MDM were detached with EDTA (5 nM); all cells were washed with PBS, incubated with Fixable Viability Stain (FVS, BD Bioscience) and washed with PBS-1% FBS. AEC cells were stained with CD95 PerCP eFluor-710 (APO-1/Fas, clone DX2) and CD178 APC (CD95L, Fas-ligand, clone NOK-1) and washed with biding buffer for annexin V PE-Cy7 stain (Thermo Scientific) and macrophages were stained with HLA-DR PE-Cy7 (clone LN3), CD163 PE (clone GHI/61), CD95 PerCP eFluor-710 (APO-1/Fas, clone DX2) and CD178 APC (CD95L, Fas-ligand, clone NOK-1). Freshly or frozen cells isolated from TA of COVID-19 patients were washed with PBS, incubated with FVS (BD Bioscience) and washed with PBS-1% FBS. Cells from TA were then stained with CD45 PE-Cy7 (clone 2D1), EpCAM PE (clone EBA-1), CD95 PerCP eFluor-710 (APO-1/Fas, clone DX2) and CD178 APC (CD95L, Fas-ligand, clone NOK-1). Lung cells from mice were isolated by proper right lung lobules digestion using collagenase (2.2 mg/mL, Sigma-Aldrich) and DNAse (0.055 mg/mL, Roche). Samples were incubated with Live/Dead viability stain (Thermo Scientific), purified anti-mouse CD16/CD32 (Fcγ III/II receptor, clone 2.4G2) and then stained with CD45 PE-Cy7 (clone 30-F11), CD31 PE-Cy7 (clone 390) and CD326 BB515 (EpCAM, clone G8.8) for AEC or stained with CD11b BV711 (clone M1/70), CD11c PE-Cy7 (clone HL3), SiglecF BB515 (clone E50-2440) and MHCII BB700 (IA/IE, clone M5/114.15.2) for macrophages, according to antibodies fabricant instructions (BD Pharmingen and eBioscience). Samples with annexin V were freshly acquired in FACS Melody (BD Biosciences) and others were fixed using PBS containing 1% paraformaldehyde (Labsynth, Diadema, SP, Brazil) and acquired in FACS Canto II (BD Biosciences). Analyses were performed in FlowJo software (Becton Dickinson and Company).

### Western blot

Calu-3 were lysed with lysis buffer and sonicated to ensure total protein release. Bradford method was used for protein dosage in the lysate supernatants (52). Samples were analyzed by SDS-PAGE and transferred onto nitrocellulose membranes. Proteins were detected using human monoclonal primary antibodies anti-cleaved caspase-3 (1:250, clone 5A1E, Cell Signaling) and anti-β-actin (1:1,000, clone 8H10D10, Cell Signaling). Primary antibodies were detected using fluorophore-conjugated secondary goat anti-rabbit (1:15,000, IRDye 680 RD, LI-COR Bioscience) or goat anti-mouse antibody (1:15,000, IRDye 800CW, LI-COR Biosciences). Fluorescent signal was detected using a LI-COR Odyssey CLx imaging system and analyzed by Image Studio software (LI-COR).

### Statistical analysis and figures

Data were analyzed using GraphPad Prism Version 8.1 (GraphPad Software, Inc., San Diego, CA, USA). Normality of data was analyzed by Shapiro-Wilk test. The comparison between two groups was performed using t-test or Mann-Whitney test. Comparisons among three groups were performed using one-way ANOVA followed by Tukey’s test or Kruskal-Wallis test. The histological score was calculated by the Chi-square test. Data were shown as the mean ± standard deviation (SD) in case of normality or shown as box with min-to-max in case of asymmetric distribution. The results were considered significant with a p-value less than 0.05. Graphics were made in GraphPad Prism and figures were made using free images from Servier Medical Art (https://smart.servier.com/). Spearman’s correlation matrix was constructed using R base functions and *qgraph* was used to build the network illustration for pair correlations.

## Acknowledgments

The authors are thankful to Ana Flávia Gembre and Denise Brufato Ferraz for excellent technical support and to Íris Castro for help with flow cytometry. This work was supported by the Sao Paulo Research Foundation (FAPESP), grants 2020/05270-0 and 2019/11213-1 to VLDB and grant 2019/09881-6 to TFCF-S, and to the Brazilian National Council for Scientific and Technological Development (CNPq), grant 312606/2019-2 to MDB.

## References

1. Sungnak W, Huang N, Bécavin C, Berg M, Queen R, Litvinukova M, et al. SARS-CoV-2 entry factors are highly expressed in nasal epithelial cells together with innate immune genes. Nat Med [Internet]. 2020 May 23;26(5):681–7. Available from: http://www.nature.com/articles/s41591-020-0868-6

2. Hou YJ, Okuda K, Edwards CE, Martinez DR, Asakura T, Dinnon KH, et al. SARS-CoV-2 Reverse Genetics Reveals a Variable Infection Gradient in the Respiratory Tract. Cell [Internet]. 2020 Jul;182(2):429-446.e14. Available from: https://linkinghub.elsevier.com/retrieve/pii/S0092867420306759

3. Grant RA, Morales-Nebreda L, Markov NS, Swaminathan S, Querrey M, Guzman ER, et al. Circuits between infected macrophages and T cells in SARS-CoV-2 pneumonia. Nature [Internet]. 2021 Feb 25;590(7847):635–41. Available from: http://www.nature.com/articles/s41586-020-03148-w

4. Zhu N, Zhang D, Wang W, Li X, Yang B, Song J, et al. A Novel Coronavirus from Patients with Pneumonia in China, 2019. N Engl J Med [Internet]. 2020 Feb 20;382(8):727–33. Available from: http://www.nejm.org/doi/10.1056/NEJMoa2001017

5. Li S, Zhang Y, Guan Z, Li H, Ye M, Chen X, et al. SARS-CoV-2 triggers inflammatory responses and cell death through caspase-8 activation. Signal Transduct Target Ther [Internet]. 2020;5(1). Available from: http://dx.doi.org/10.1038/s41392-020-00334-0

6. Blanco-Melo D, Nilsson-Payant BE, Liu W-C, Uhl S, Hoagland D, Møller R, et al. Imbalanced Host Response to SARS-CoV-2 Drives Development of COVID-19. Cell [Internet]. 2020 May;181(5):1036-1045.e9. Available from: https://linkinghub.elsevier.com/retrieve/pii/S009286742030489X

7. Dorward DA, Russell CD, Um IH, Elshani M, Armstrong SD, Penrice-Randal R, et al. Tissue-specific Immunopathology in Fatal COVID-19. Am J Respir Crit Care Med [Internet]. 2020 Nov 20;rccm.202008-3265OC. Available from: https://www.atsjournals.org/doi/10.1164/rccm.202008-3265OC

8. Fan E, Beitler JR, Brochard L, Calfee CS, Ferguson ND, Slutsky AS, et al. COVID-19-associated acute respiratory distress syndrome: is a different approach to management warranted? Lancet Respir Med [Internet]. 2020 Aug;8(8):816–21. Available from: https://linkinghub.elsevier.com/retrieve/pii/S2213260020303040

9. Menter T, Haslbauer JD, Nienhold R, Savic S, Hopfer H, Deigendesch N, et al. Postmortem examination of COVID-19 patients reveals diffuse alveolar damage with severe capillary congestion and variegated findings in lungs and other organs suggesting vascular dysfunction. Histopathology [Internet]. 2020 Aug 5;77(2):198–209. Available from: https://onlinelibrary.wiley.com/doi/10.1111/his.14134

10. Ziegler CGK, Allon SJ, Nyquist SK, Mbano IM, Miao VN, Tzouanas CN, et al. SARS-CoV-2 Receptor ACE2 Is an Interferon-Stimulated Gene in Human Airway Epithelial Cells and Is Detected in Specific Cell Subsets across Tissues. Cell [Internet]. 2020;181(5):1016-1035.e19. Available from: http://www.ncbi.nlm.nih.gov/pubmed/32413319

11. Liu F, Han K, Blair R, Kenst K, Qin Z, Upcin B, et al. SARS-CoV-2 Infects Endothelial Cells In Vivo and In Vitro. Front Cell Infect Microbiol [Internet]. 2021 Jul 6;11. Available from: https://www.frontiersin.org/articles/10.3389/fcimb.2021.701278/full

12. Varga Z, Flammer AJ, Steiger P, Haberecker M, Andermatt R, Zinkernagel AS, et al. Endothelial cell infection and endotheliitis in COVID-19. Lancet [Internet]. 2020 May;395(10234):1417–8. Available from: https://linkinghub.elsevier.com/retrieve/pii/S014067362030917X

13. Mehta P, McAuley DF, Brown M, Sanchez E, Tattersall RS, Manson JJ. COVID-19: consider cytokine storm syndromes and immunosuppression. Lancet [Internet]. 2020;395(10229):1033–4. Available from: http://dx.doi.org/10.1016/S0140-6736(20)30628-0

14. Dewaele K, Claeys R. Hemophagocytic lymphohistiocytosis in SARS-CoV-2 infection. Blood [Internet]. 2020 Jun 18;135(25):2323–2323. Available from: https://ashpublications.org/blood/article/135/25/2323/460907/Hemophagocytic-lymphohistiocytosis-in-SARSCoV2

15. Fraga-Silva TF de C, Maruyama SR, Sorgi CA, Russo EM de S, Fernandes APM, de Barros Cardoso CR, et al. COVID-19: Integrating the Complexity of Systemic and Pulmonary Immunopathology to Identify Biomarkers for Different Outcomes. Front Immunol [Internet]. 2021 Jan 29;11. Available from: https://www.frontiersin.org/articles/10.3389/fimmu.2020.599736/full

16. Mahmoodpoor A, Hosseini M, Soltani-Zangbar S, Sanaie S, Aghebati-Maleki L, Saghaleini SH, et al. Reduction and exhausted features of T lymphocytes under serological changes, and prognostic factors in COVID-19 progression. Mol Immunol [Internet]. 2021 Oct;138:121–7. Available from: https://linkinghub.elsevier.com/retrieve/pii/S0161589021001760

17. Nienhold R, Ciani Y, Koelzer VH, Tzankov A, Haslbauer JD, Menter T, et al. Two distinct immunopathological profiles in autopsy lungs of COVID-19. Nat Commun [Internet]. 2020 Dec 8;11(1):5086. Available from: https://www.nature.com/articles/s41467-020-18854-2

18. Desai N, Neyaz A, Szabolcs A, Shih AR, Chen JH, Thapar V, et al. Temporal and spatial heterogeneity of host response to SARS-CoV-2 pulmonary infection. Nat Commun [Internet]. 2020 Dec 9;11(1):6319. Available from: http://www.nature.com/articles/s41467-020-20139-7

19. Veras FP, Pontelli M, Silva C, Toller-Kawahisa J, de Lima M, Nascimento D, et al. SARS-CoV-2 triggered neutrophil extracellular traps (NETs) mediate COVID-19 pathology. medRxiv [Internet]. 2020; Available from: https://www.medrxiv.org/content/10.1101/2020.06.08.20125823v1

20. Giamarellos-Bourboulis EJ, Netea MG, Rovina N, Akinosoglou K, Antoniadou A, Antonakos N, et al. Complex Immune Dysregulation in COVID-19 Patients with Severe Respiratory Failure. Cell Host Microbe [Internet]. 2020 Jun;27(6):992-1000.e3. Available from: https://linkinghub.elsevier.com/retrieve/pii/S1931312820302365

21. Ulhaq ZS, Soraya GV. Interleukin-6 as a potential biomarker of COVID-19 progression. Médecine Mal Infect [Internet]. 2020 Jun;50(4):382–3. Available from: https://linkinghub.elsevier.com/retrieve/pii/S0399077X20300883

22. Mojtabavi H, Saghazadeh A, Rezaei N. Interleukin-6 and severe COVID-19: a systematic review and meta-analysis. Eur Cytokine Netw [Internet]. 2020 Jun;31(2):44–9. Available from: http://www.john-libbey-eurotext.fr/medline.md?doi=10.1684/ecn.2020.0448

23. Gorham J, Moreau A, Corazza F, Peluso L, Ponthieux F, Talamonti M, et al. Interleukine-6 in critically ill COVID-19 patients: A retrospective analysis. Lazzeri C, editor. PLoS One [Internet]. 2020 Dec 31;15(12):e0244628. Available from: https://dx.plos.org/10.1371/journal.pone.0244628

24. Fumagalli MJ, Castro-Jorge LA, Fraga-Silva TF de C, de Azevedo PO, Capato CF, Rattis BAC, et al. Protective Immunity against Gamma and Zeta Variants after Inactivated SARS-CoV-2 Virus Immunization. Viruses [Internet]. 2021 Dec 4;13(12):2440. Available from: https://www.mdpi.com/1999-4915/13/12/2440

25. Chu H, Chan JF-W, Yuen TT-T, Shuai H, Yuan S, Wang Y, et al. Comparative tropism, replication kinetics, and cell damage profiling of SARS-CoV-2 and SARS-CoV with implications for clinical manifestations, transmissibility, and laboratory studies of COVID-19: an observational study. The Lancet Microbe [Internet]. 2020 May;1(1):e14–23. Available from: https://linkinghub.elsevier.com/retrieve/pii/S2666524720300045

26. Zhu N, Wang W, Liu Z, Liang C, Wang W, Ye F, et al. Morphogenesis and cytopathic effect of SARS-CoV-2 infection in human airway epithelial cells. Nat Commun [Internet]. 2020 Dec 6;11(1):3910. Available from: https://www.nature.com/articles/s41467-020-17796-z

27. Liu Y, Garron TM, Chang Q, Su Z, Zhou C, Qiu Y, et al. Cell-Type Apoptosis in Lung during SARS-CoV-2 Infection. Pathogens [Internet]. 2021 Apr 23;10(5):509. Available from: https://www.mdpi.com/2076-0817/10/5/509

28. Ren Y, Shu T, Wu D, Mu J, Wang C, Huang M, et al. The ORF3a protein of SARS-CoV-2 induces apoptosis in cells. Cell Mol Immunol [Internet]. 2020 Aug 18;17(8):881–3. Available from: http://www.nature.com/articles/s41423-020-0485-9

29. Ampomah PB, Lim LHK. Influenza A virus-induced apoptosis and virus propagation. Apoptosis [Internet]. 2020 Feb;25(1–2):1–11. Available from: http://link.springer.com/10.1007/s10495-019-01575-3

30. O’Donnell DR, Milligan L, Stark JM. Induction of CD95 (Fas) and Apoptosis in Respiratory Epithelial Cell Cultures Following Respiratory Syncytial Virus Infection. Virology [Internet]. 1999 Apr;257(1):198–207. Available from: https://linkinghub.elsevier.com/retrieve/pii/S0042682299996502

31. Fujimoto I, Takizawa T, Ohba Y, Nakanishi Y. Co-expression of Fas and Fas-ligand on the surface of influenza virus-infected cells. Cell Death Differ [Internet]. 1998 May 15;5(5):426–31. Available from: http://www.nature.com/articles/4400362

32. Nienhold R, Ciani Y, Koelzer VH, Tzankov A, Haslbauer JD, Menter T, et al. Two distinct immunopathological profiles in autopsy lungs of COVID-19. Nat Commun [Internet]. 2020;11(1):1–13. Available from: http://dx.doi.org/10.1038/s41467-020-18854-2

33. Hagimoto N, Kuwano K, Kawasaki M, Yoshimi M, Kaneko Y, Kunitake R, et al. Induction of Interleukin-8 Secretion and Apoptosis in Bronchiolar Epithelial Cells by Fas Ligation. Am J Respir Cell Mol Biol [Internet]. 1999 Sep;21(3):436–45. Available from: http://www.atsjournals.org/doi/abs/10.1165/ajrcmb.21.3.3397

34. Cullen SP, Henry CM, Kearney CJ, Logue SE, Feoktistova M, Tynan GA, et al. Fas/CD95-Induced Chemokines Can Serve as “Find-Me” Signals for Apoptotic Cells. Mol Cell [Internet]. 2013 Mar;49(6):1034–48. Available from: https://linkinghub.elsevier.com/retrieve/pii/S1097276513000890

35. Choi C, Gillespie GY, Van Wagoner NJ, Benveniste EN. Fas engagement increases expression of interleukin-6 in human glioma cells. J Neurooncol [Internet]. 2002 Jan;56(1):13–9. Available from: http://www.ncbi.nlm.nih.gov/pubmed/11949822

36. Liu X, Wang H, Shi S, Xiao J. Association between IL-6 and severe disease and mortality in COVID-19 disease: a systematic review and meta-analysis. Postgrad Med J [Internet]. 2021 Jun 3;postgradmedj-2021-139939. Available from: https://pmj.bmj.com/lookup/doi/10.1136/postgradmedj-2021-139939

37. Peter ME, Budd RC, Desbarats J, Hedrick SM, Hueber A-O, Newell MK, et al. The CD95 Receptor: Apoptosis Revisited. Cell [Internet]. 2007 May;129(3):447–50. Available from: https://linkinghub.elsevier.com/retrieve/pii/S0092867407005429

38. Guégan JP, Ginestier C, Charafe-Jauffret E, Ducret T, Quignard J-F, Vacher P, et al. CD95/Fas and metastatic disease: What does not kill you makes you stronger. Semin Cancer Biol [Internet]. 2020 Feb;60:121–31. Available from: https://linkinghub.elsevier.com/retrieve/pii/S1044579X19301294

39. Park DR, Thomsen AR, Frevert CW, Pham U, Skerrett SJ, Kiener PA, et al. Fas (CD95) Induces Proinflammatory Cytokine Responses by Human Monocytes and Monocyte-Derived Macrophages. J Immunol [Internet]. 2003 Jun 15;170(12):6209–16. Available from: http://www.jimmunol.org/lookup/doi/10.4049/jimmunol.170.12.6209

40. Altemeier WA, Zhu X, Berrington WR, Harlan JM, Liles WC. Fas (CD95) induces macrophage proinflammatory chemokine production via a MyD88-dependent, caspase-independent pathway. J Leukoc Biol [Internet]. 2007 Sep;82(3):721–8. Available from: http://doi.wiley.com/10.1189/jlb.1006652

41. Melms JC, Biermann J, Huang H, Wang Y, Nair A, Tagore S, et al. A molecular single-cell lung atlas of lethal COVID-19. Nature [Internet]. 2021 Jul 1;595(7865):114–9. Available from: https://www.nature.com/articles/s41586-021-03569-1

42. Jones SA, Hunter CA. Is IL-6 a key cytokine target for therapy in COVID-19? Nat Rev Immunol [Internet]. 2021 Jun 13;21(6):337–9. Available from: http://www.nature.com/articles/s41577-021-00553-8

43. Elahi R, Karami P, Heidary AH, Esmaeilzadeh A. An updated overview of recent advances, challenges, and clinical considerations of IL-6 signaling blockade in severe coronavirus disease 2019 (COVID-19). Int Immunopharmacol [Internet]. 2022 Apr;105:108536. Available from: https://linkinghub.elsevier.com/retrieve/pii/S1567576922000200

44. Canziani LM, Trovati S, Brunetta E, Testa A, De Santis M, Bombardieri E, et al. Interleukin-6 receptor blocking with intravenous tocilizumab in COVID-19 severe acute respiratory distress syndrome: A retrospective case-control survival analysis of 128 patients. J Autoimmun [Internet]. 2020 Nov;114:102511. Available from: https://linkinghub.elsevier.com/retrieve/pii/S0896841120301335

45. Kewan T, Covut F, Al–Jaghbeer MJ, Rose L, Gopalakrishna KV, Akbik B. Tocilizumab for treatment of patients with severe COVID–19: A retrospective cohort study. EClinicalMedicine [Internet]. 2020 Jul;24:100418. Available from: https://linkinghub.elsevier.com/retrieve/pii/S2589537020301620

46. Kaye AG, Siegel R. The efficacy of IL-6 inhibitor Tocilizumab in reducing severe COVID-19 mortality: a systematic review. PeerJ [Internet]. 2020 Nov 2;8:e10322. Available from: https://peerj.com/articles/10322

47. Araujo DB, Machado RRG, Amgarten DE, Malta F de M, de Araujo GG, Monteiro CO, et al. SARS-CoV-2 isolation from the first reported patients in Brazil and establishment of a coordinated task network. Mem Inst Oswaldo Cruz [Internet]. 2020;115. Available from: http://www.scielo.br/scielo.php?script=sci_arttext&pid=S0074-02762020000100344&tlng=en

48. Baxter EW, Graham AE, Re NA, Carr IM, Robinson JI, Mackie SL, et al. Standardized protocols for differentiation of THP-1 cells to macrophages with distinct M(IFNγ+LPS), M(IL-4) and M(IL-10) phenotypes. J Immunol Methods. 2020;478(December):1–11.

49. Hoepel W, Chen H-J, Geyer CE, Allahverdiyeva S, Manz XD, de Taeye SW, et al. High titers and low fucosylation of early human anti–SARS-CoV-2 IgG promote inflammation by alveolar macrophages. Sci Transl Med [Internet]. 2021 Jun 2;13(596). Available from: https://www.science.org/doi/10.1126/scitranslmed.abf8654

50. Edgar R. Gene Expression Omnibus: NCBI gene expression and hybridization array data repository. Nucleic Acids Res [Internet]. 2002 Jan 1;30(1):207–10. Available from: https://academic.oup.com/nar/article-lookup/doi/10.1093/nar/30.1.207

51. Benjamini Y, Hochberg Y. Controlling the False Discovery Rate: A Practical and Powerful Approach to Multiple Testing. J R Stat Soc Ser B [Internet]. 1995 Mar 10;57(1):289–300. Available from: http://www.jstor.org/stable/2346101

52. Bradford M. A Rapid and Sensitive Method for the Quantitation of Microgram Quantities of Protein Utilizing the Principle of Protein-Dye Binding. Anal Biochem [Internet]. 1976 May 7;72(1– 2):248–54. Available from: http://linkinghub.elsevier.com/retrieve/pii/S0003269776699996

